# Dissecting the transcriptional regulation of infant and childhood acute myeloid leukemia

**DOI:** 10.1101/2024.08.14.608016

**Authors:** Rasoul Godini

## Abstract

Pediatric acute myeloid leukemia (AML) exhibits distinct characteristics between infants and children, manifested by variations in clinical features, cytogenetic abnormalities, and molecular aberrations. While most studies have focused on the associations between genomic abnormalities, age groups, and disease status, the transcriptional profiles and gene regulatory mechanisms underlying these differences remain underexplored. In this study, through differential gene expression analysis and Weighted Gene Co-expression Network Analysis, I have identified gene groups associated with AML and further categorized them by infant and childhood subgroups. The findings reveal three distinct gene groups with age- and tissue-specific expression patterns. Additionally, I propose a gene regulatory circuitry that elucidates the differences between infant and childhood AML. Several novel markers demonstrating significant expression changes in tumors were identified. Moreover, a comprehensive gene regulatory network for pediatric AML was constructed using differentially expressed transcription factors and protein-gene interaction data. This study highlights new gene regulation mechanisms in pediatric AML and offers potential avenues for developing novel biomarkers and therapeutic strategies.

## Introduction

Cancer is the foremost cause of mortality in individuals under 19 years of age in the United States [1]. According to US Cancer Statistics spanning from 2003 to 2019, leukemia exhibits the highest incidence rate among pediatric malignancies, at 46.6 per 1 million, with acute myeloid leukemia (AML) comprising approximately 16.09% of these cases [2]. Pediatric leukemia show minimal sex disparities, with a higher incidence rate in male compared to female (1.27, 95% CI = 1.24 to 1.29), Nonetheless, no statistically significant difference is evident in pediatric AML (1.02, 95% CI = 0.99 to 1.06) [2]. A notable characteristic of pediatric AML is its heterogeneity in morphology, cytogenetics, immunophenotype, genetic abnormalities, and clinical behaviour [3–5]. These characteristics are used, usually in combination, to classify the diseases and select treatment approaches [3]. A prominent distinction is observed in infant AML patients (age < 2 years) compared to other pediatric patients that manifest unfavourable risk factors and higher susceptibility to therapy-associated toxicity [6–9]. Despite the challenges associated with infant AML group, the treatment strategies for these patients have been improved [10]. Nonetheless, a comprehensive characterization distinguishing infant from childhood AML could further enhance treatment approaches.

The heterogeneity in pediatric AML is classified into various categories: clinical features, cytogenetic abnormalities, and molecular aberrations. Infant AML patients exhibit a higher incidence of acute monoblastic, megakaryoblastic, and myelomonoblastic leukemia. Additionally, they present with increased involvement of the central nervous system and extramedullary organs [11, 12]. Infants with AML manifest a distinct cytogenic abnormalities compared to other pediatric AML patients [11]. For example, in infant AML patients, translocations frequently involve chromosome band 11q23, whereas core-binding factor abnormalities and t(15;17), which are favorable cytogenetic features, are less common [11, 13]. Furthermore, KMT2A (MLL1) rearrangement is more prevalent (∼14%) in infants with AML, decreasing to 4-7% in older patients [11, 14]. Distinct features in mutations and molecular pathway aberrations are also observed in infant AML patients. Unlike rearrangements, KMT2A (MLL1) mutations are rare in children and undetected in infant AML patients [14]. Mutations in nucleophosmin 1, FLT3 and CEBPA genes are also rare in infant with AML [15–18]. The CBFA2T3-GLIS2 fusion transcript is a more frequent abnormality in infants compared to children [19]. Generally, a similar frequency of mutations in RAS pathway has been observed in infants and children with AML, with a comparable mutation rate in N-RAS and PTPN11 genes and a higher rate in children over 2 years for K-RAS gene [14, 20]. The application of genomic techniques that provide high-throughput data has unveiled new insights, further distinguishing pediatric AML from adult AML [21, 22].

Genomic data has facilitated the discovery of subtypes of pediatric AML, enhancing the selection of appropriate treatment strategies [5]. Despite progresses in pediatric AML prognosis and treatment, there is a lack of knowledge in physiological and regulatory mechanisms distinguishing infants from older childhood AML patients that yet need to be elucidated. In this study, I aim to identify the transcriptional differences between infants and children with AML that confer these distinctions. I compared two populations of patients—infants (0-2 years) and children (3-19 years)—regardless of genomic abnormalities, to obtain a comprehensive landscape of gene alterations between these age groups. I utilize the Therapeutically Applicable Research to Generate Effective Treatments (TARGET) database, which provides a large cohort of pediatric AML genomic data, to investigate the transcriptome and gene regulatory mechanisms associated with infant and childhood AML [23]. Many studies consider age < 2 as the infants due to similar physiological characteristics and treatment outcomes [8, 9, 24]. Therefore, in this study, I classify patients under 2 years of age as infant AML cases. My results identify several genes associated with the distinctions between infant and childhood AML. I construct regulatory networks of transcription factors (TFs) that potentially regulate pediatric AML and the differences between infants and older patients. These findings can improve age-specific diagnostic and treatment strategies for pediatric AML and contribute to the discovery of new therapeutic mechanisms.

## Methods

### Data preparation

Transcriptome profiling, methylation, and Copy Number Variation (CNV) data were obtained from the Genomic Data Commons (GDC) through the TARGET project [23], using TCGAbiolinks package in the R programming language [25]. All datasets were filtered to include bone marrow samples paired with healthy tissues (collected from the same individuals) from patients aged 0-19 years. A total of 406 samples (203 tumor and 203 normal) were used for transcriptome analysis. Using paired samples for comparisons helps to mitigate unwanted sources of batch effects. For methylation analysis, 164 samples (82 tumor and 82 normal) were used, and for CNV analysis, 49 samples (all tumor) were included. In the majority of cases, transcriptome, methylation, and CNV data were obtained from different individuals, thus using different patients for the analyses was unavoidable.

### Differentially expressed gene analysis

To perform differentially expressed analysis on multiple samples Wilcoxon signed rank test (paired samples) or Wilcoxon rank-sum test (unpaired samples) was applied [26]. This method has the advantage of more robust P-values as the popular methods, such as edgeR and DESeq2 result in large numbers of false positive Differentially Expressed Genes (DEGs) [26]. Prior to DEG analysis, samples were filtered for genes with adequate expression levels, normalized for library sizes, and log-transformed using edgeR package in R [26]. If further filtering requirements (age/sex/age group) samples were selected accordingly. The low expressed genes were excluded using “filterByExpr” function from edgeR package in R [27], if they had a read count < 10 in less than 75% of samples of the smallest group (tumor/normal) and a total read count of < 200 across all samples. Following this, the trimmed mean of M-values (TMM) normalization was performed, and counts per million (CPM) were produced using the edgeR package in R [27]. For differentially expression analysis the CPM, and for visualising and WGCNA the log2CPM values were used [26]. Genes were identified as DEGs if they met the following thresholds: False Discovery Rate (FDR) <= 0.05, and log2 Fold Change (FC) ± 1 Log2FC or ± 1.5 Log2FC.

### Differentially methylated region analysis

For methylation analysis, methylation beta values obtained from the Illumina Human Methylation 450 platform were utilized. The data were refined by removing unreliable probes through the following filters: (i) probes containing small insertions and deletions (INDELs), repetitive DNA, single nucleotide polymorphisms (SNPs), and regions of reduced genomic complexity [28], and (ii) cross-reactive probes [29]. Lowly expressed probes were omitted using the “filterByExpr” function from the edgeR package, with the following thresholds: min.count = 0.02, min.prop = 0.85, and min.total.count = 10 [27]. All other probes with low expression, as well as those located in different genomic regions (islands, shores, and open seas), were retained for further analysis. Differentially Methylated Regions (DMRs) were identified using the Wilcoxon signed-rank test [26] and DMRs were selected based on an FDR < 0.05, a log2 fold change of ±0.378 (corresponding to a 1.2 change in beta value), and their association with transcription start sites (within 100 or 1500 bp of the TSS), gene bodies, and other regions (within 5’UTR, 1st Exon, and 3’UTR).

### Copy Number Variation

For CNV analysis I applied public free accessible “Gene Level Copy Number” data type from TARGET. Data was filtered for bone marrow samples of patients aged < 20. The median copy number was used as CNV value.

### Weighted gene co-expression network analysis

To construct a network of co-expressed genes associated with biological states, the Weighted Gene Co-expression Network Analysis (WGCNA) method was employed [30]. Samples were categorized into normal and tumor groups, divided into age groups 0-2 years and 3-19 years, as well as an all-ages group (0-19 years), in addition to sexes. Following the removal of lowly expressed genes (as described in the Differentially Expressed Gene analysis section), the top 20,000 most variable genes were selected and analyzed using the WGCNA package in R. A “signed” network was constructed with a soft-power threshold of 16 (Supp. Fig 2). The following parameters were applied to select modules: “hybrid” method, deep split = 2, and module size > 50. The modules most highly associated with the biological states were chosen for further analysis. Module visualization and analysis were performed using Cytoscape 3.10.1 [31]. In each network, weighted degree and betweenness values were utilized to identify central nodes [32].

### Gene ontology enrichment analysis

Gene annotation analysis was performed using Database for Annotation, Visualization and Integrated Discovery (DAVID) version (v2023q4) [33]. I only used top 10 Biological Processes (BP) with a P-value < 0.05.

### Gene regulatory network construction

Transcriptional regulatory networks were constructed by mapping the Transcription Factors (TFs) of interest onto the hTFtarget network, obtained from “guolab.wchscu.cn/hTFtarget/” [34]. hTFtarget encompasses interactions between TFs and their targets based on chromatin immunoprecipitation sequencing (ChIP-seq) data from 7,190 experiments across various cell lines and tissues. The list of human TFs (1,639 TFs) provided by Lambert and colleagues was used as the reference for selecting TFs [35]. Briefly, the interaction map of the TFs identified through differential expression and WGCNA analyses was extracted from the extensive list of interactions (1,342,129 interactions) between TFs and their corresponding targets [34]. The resulting interactions were filtered to include only those involving TFs identified in bone marrow tissue. The list of oncogenes was obtained from “cancer.sanger.ac.uk” [36] and assigned to the relevant TFs. The networks were visualized using Cytoscape 3.10.1 [31].

## Results

### Experimental design

I aimed to uncover age- and sex-associated gene expression heterogeneity in pediatric AML patients. To achieve this, I utilized the TARGET database to establish a robust cohort by filtering for primary tumor samples collected from the bone marrow of patients under 20 years of age. Additionally, I included only paired normal and tumor samples from the same patients to minimize inter-patient variation. Two approaches were employed for sample analysis: (i) an unbiased approach to compare age groups in 5-year intervals (0-4, 5-9, 10-14, 15-19), which allows for comparison across age groups without regard to infant and childhood phenotypes, and (ii) a comparative analysis between infants (0-2) and children (3-19) using DEGs and WGCNA (Fig. 1).

**Figure 1.**
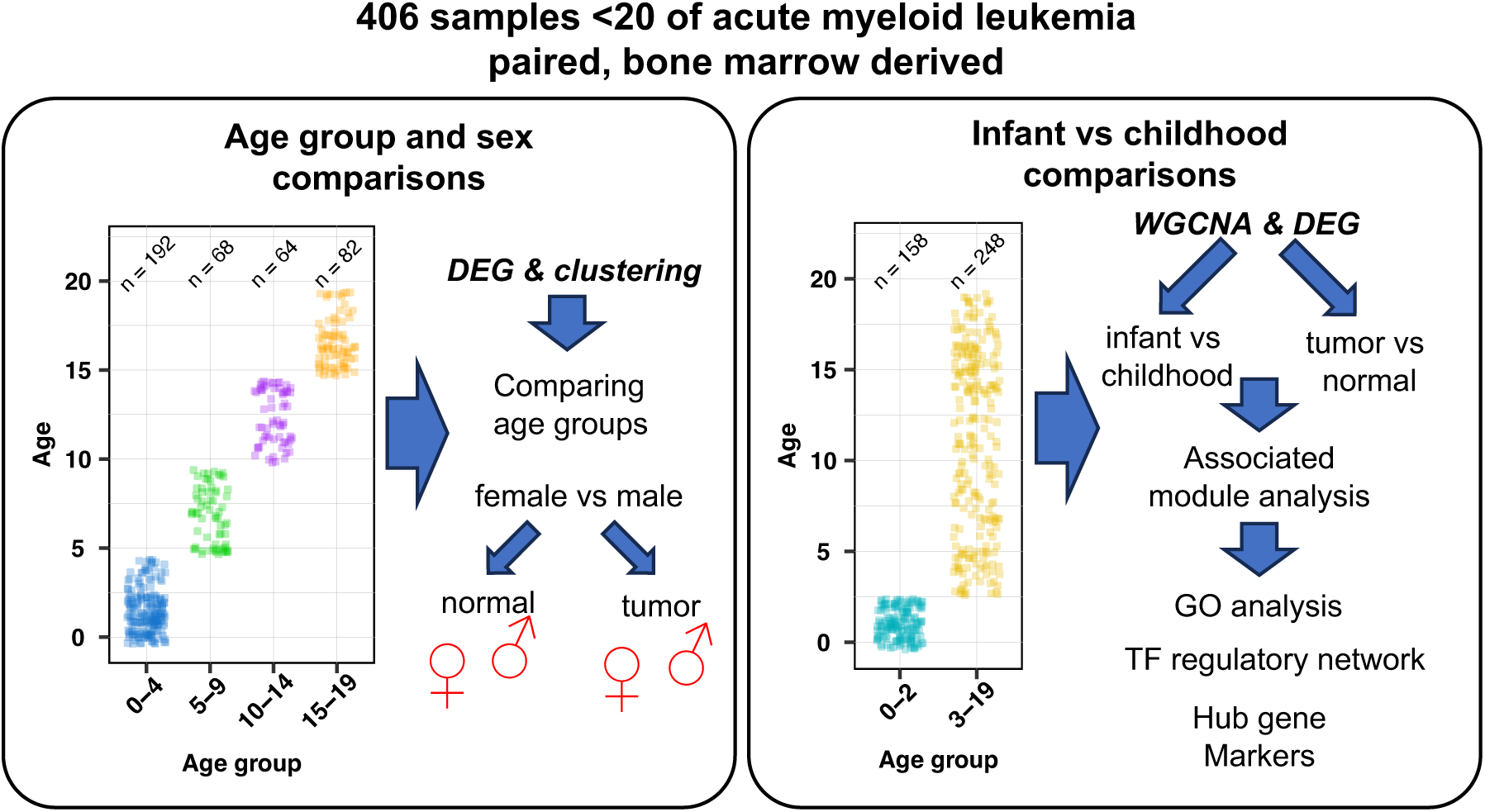
A general overview of the analyses conducted. To ensure a homogeneous population, the original TARGET dataset was filtered for primary blood-derived tumor samples from bone marrow paired with normal tissue samples from the same patient. Additionally, samples were filtered based on ages <20. **Left side)** unbiased combinatorial comparisons were performed between age groups, different sexes, and tissue types to identify potential differences among subgroups. **Right side)** samples were sorted into two age groups: 0-2 as infant and 3-19 as childhood samples. Weighted Gene Co-expression Network Analysis (WGCNA) and Differential Expressed Genes (DEG) analyses were performed to identify gene expression patterns associated with each age group and tumor samples. Transcriptional regulatory networks controlling age groups and tumors were constructed, and hub Transcription Factors (TFs) were identified. Finally, several new potential markers are introduced based on graph theory principles and differential expression categories.

In the unbiased approach, I examined both age and sex-associated differences to identify potential sex disparities. For the infant and childhood analyses, sex differences were compared between the two age groups within each tissue type to identify age- and tissue-specific changes between sexes. Furthermore, I sought age- and tissue-associated gene signatures by applying WGCNA and constructed a potentially regulatory network to differentiate between infant and childhood AML. Finally, I used DEGs and WGCNA results to identify novel markers and regulatory networks in pediatric AML.

### Age and sex associated gene signature in pediatric AML

To identify potential sex-specific gene signatures in pediatric AML, I investigated sex-associated genes by examining highly variable genes between sexes across all samples and different age groups. Comparison of variation among samples using the top 3000 most variable genes indicated a clear separation between normal and tumor samples but did not reveal any distinction based on sex (Fig. 2A). However, Principal Component Analysis (PCA) of the top 100 most variable genes demonstrated clustering of samples according to sex (Fig. 2A). This sex-specific clustering is attributable to Y chromosome-associated genes, as removing Y chromosome transcripts from the transcriptome eliminated this clustering (Supp. Fig. 1 A). I then performed PCA using the top 50 most variable genes, focusing specifically on sex and tissue types to identify age-associated clusters. This approach removed the effects of sex and tissue, allowing for an exclusive analysis of age groups (Fig. 2A and Supp. Fig. 1 B). The results revealed no distinct clustering of age-associated samples. Nonetheless, a subtle clustering within the 0-4 age group was observed among male tumor samples, distinguishing them from other samples (Fig. 1A).

**Figure 2.**
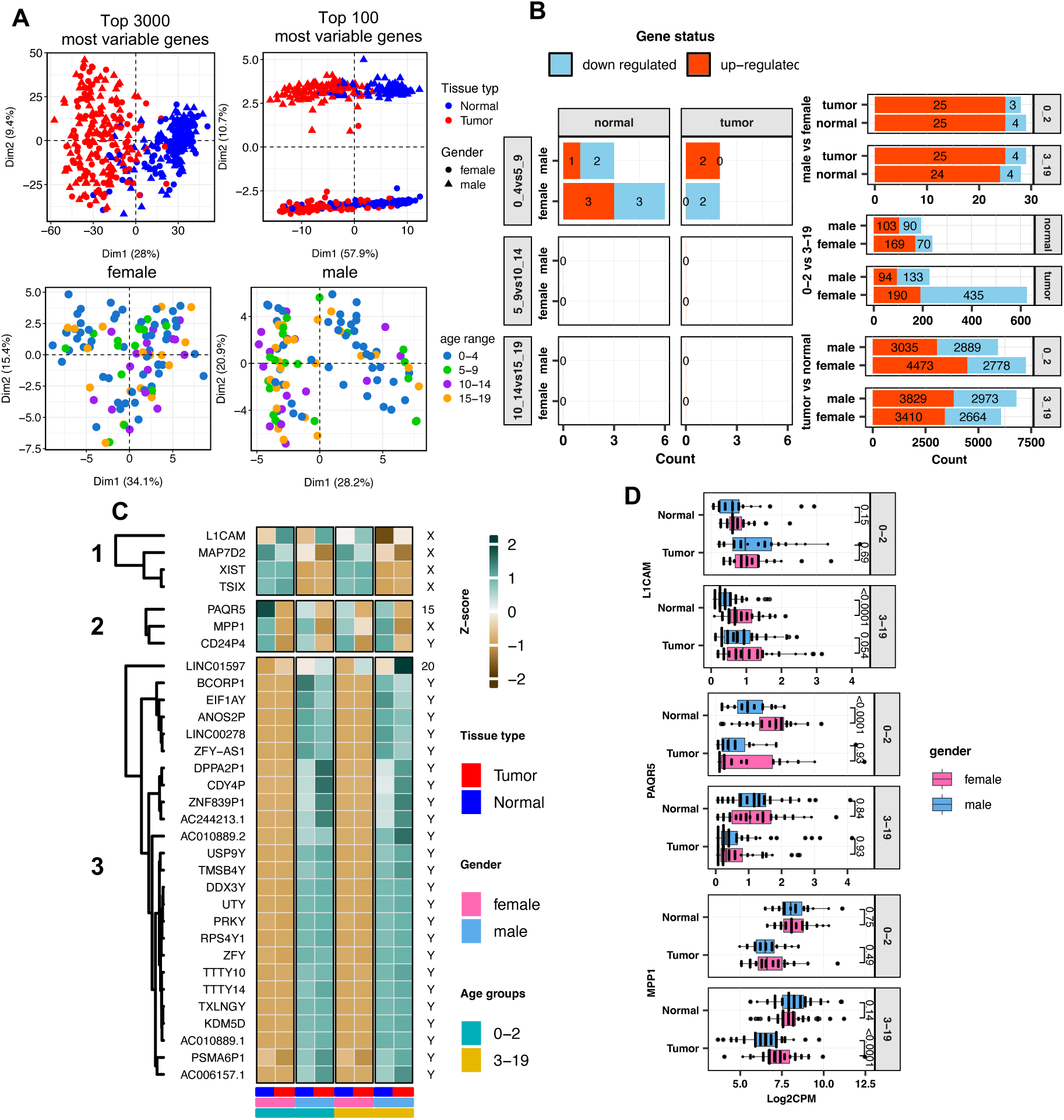
Unbiased comparisons of age groups, sexes and tissue types. **A)** PCA of the most variable genes in each sample. Top plots show all 406 samples clustered based on the tissue type and sex. Two sets of 100 and 3000 most variable genes were used to identify potential clustering capacities in sexes and tissue types. Bottom plots show clustering of age groups in each sex. **B)** Number of DEGs within and between age groups, sexes and tissues. The left plot shows unbiased comparison between samples aged < 20 in five years bins to identify age-associated DEGs. Right plots represent combinatorial comparisons between 0-2 and 3-19 and different sexes and tissue types. For example, the top right plot, DEGs is the result of comparing tumor tissues in male and female samples. **C)** Heatmap showing expression of DEGs between female and male. The heatmap columns represent median of expression in different combinations of age, sex and tissue type. **D)** Bar charts compare the expression of candidate non-Y chromosome DEGs between female and male.

I also identified DEGs across various tissues, age groups, and sexes (Fig. 1B and Supp. file 1 S1-S6). When comparing all tumor versus normal samples, while ignoring age and sex, 2650 genes were found to be down-regulated and 3744 genes were up-regulated (Supp. Fig. 1C and Supp. file 1 S1). In contrast, comparison of all male versus female samples, ignoring tissue and age, identified 25 up-regulated genes and 3 down-regulated genes (Supp. Fig. 1C and Supp. file 1 S1). Analysis of DEGs between age subgroups revealed a few differentially expressed genes between the 0-4 and 5-9 age groups in both tumor and normal tissues for male and female patients. No DEGs were observed in comparisons of other age groups (Fig. 2B, left plot). Based on these observations, I focused on comparing infant (0-2) and childhood (3-19) AML, which overlap with the age groups of 0-4 and 5-9. To control for potential confounders, multiple comparisons were conducted using a combinatorial approach that included comparisons based on sex, age, and tissue type. For instance, female and male samples were compared within the 0-2 and 3-19 age groups for both normal and tumor samples. The results indicated that the fewest DEGs were observed between female and male samples, followed by comparisons between age groups, with the highest number of DEGs found in tissue comparisons (Fig. 2B, right plot). To investigate sex-associated genes within each age group and tissue type, I clustered all DEGs identified from comparisons between male and female samples in different combinations (Fig. 2B) using a heatmap to identify non-Y chromosome genes exhibiting sex-specific expression patterns (Fig. 2C).

I identified four genes (L1CAM, PAQR5, LINC01597, and MPP1) that are associated with sex. Among these, LINC01597 exhibited relatively low expression levels across all samples (Supp. Fig. 1D). Consequently, I focused further analysis on L1CAM, MPP1, and PAQR5. L1CAM and PAQR5 were found to be expressed at higher levels in normal tissues from females compared to males, specifically in the infant (0-2) and childhood (3-19) age groups, respectively (Fig. 2D). MPP1 showed increased expression in female tumor samples from childhood patients (Fig. 2D). For additional analysis, due to the very limited number of sex-associated genes, I combined the female and male samples to enhance the statistical robustness of the analysis.

Overall, the results indicate that transcriptomic differences in pediatric AML patients are primarily associated with the transition from infancy to childhood. Moreover, at least three genes are differentially expressed between female and male patients in a tissue- and age-specific manner.

### WGCNA identifies gene groups related to age-associated heterogeneity of pediatric AML

Identifying genes associated with infant and childhood AML requires uncovering gene expressions that are specific to age and tissue. This means the gene expression should remain independent of changes in the counterpart tissue as the patient ages. Conventional PCA, analysis of all samples, and separate analyses of normal or tumor samples failed to cluster the samples according to age (Fig. 3A). Additionally, hierarchical clustering did not group samples based on age categories (Fig. 3B). To address this, two approaches were employed to identify age-specific genes: (i) identifying DEGs that are age- and tissue-specific, meaning that expression changes should be observed between age-specific subgroups of samples, and (ii) genes associated to age-groups of each tissue types identified by WGCNA. These approaches yielded a list of genes suitable for further analysis.

**Figure 3.**
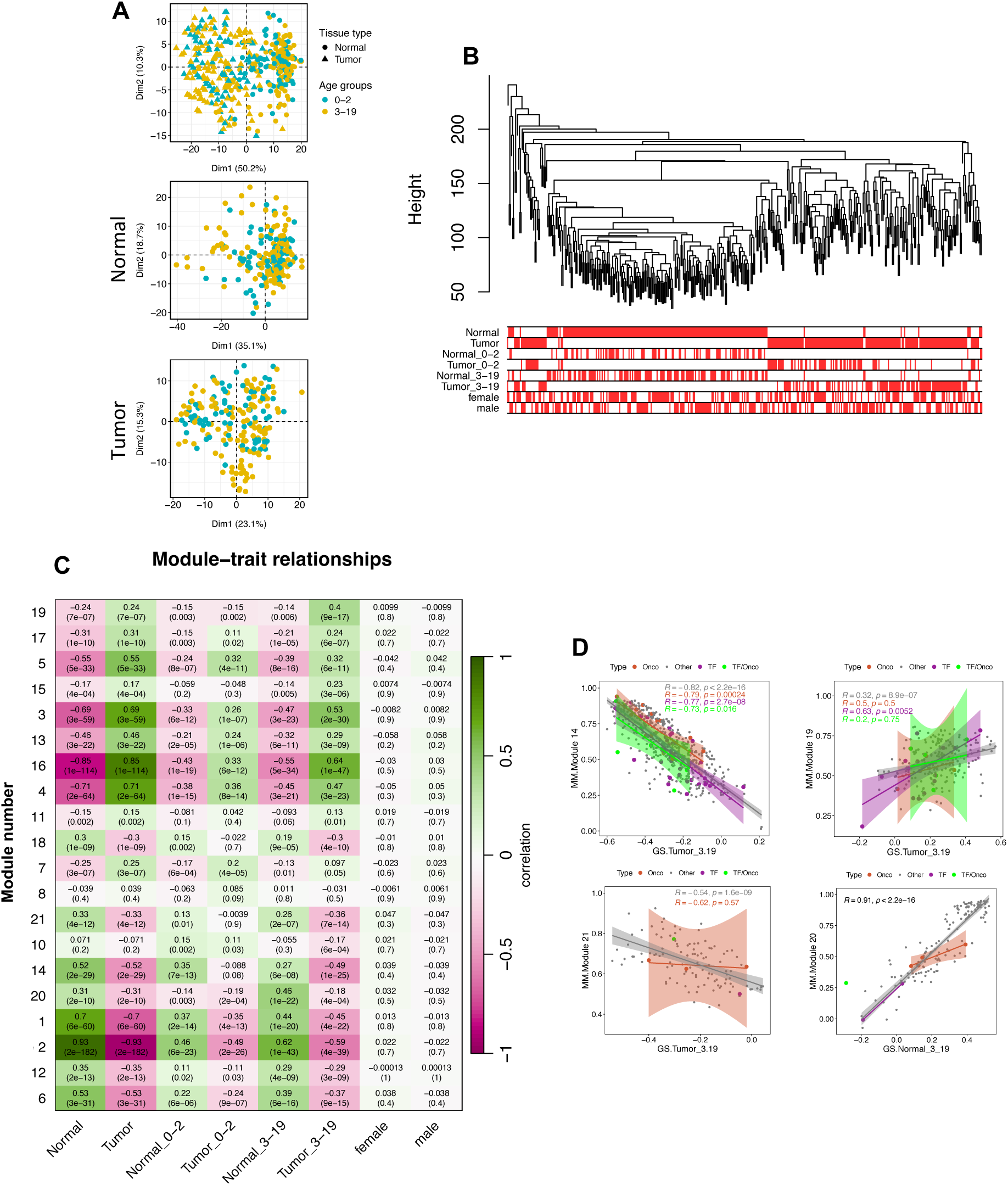
Weighted Gene Co-expression Network Analysis (WGCNA) of paired samples. **A)** PCA analysis of samples based on age groups in tumor, normal and both samples. PCA is performed using top 300 most variable genes. **B)** Hierarchical clustering of samples based on tissue, age-tissue and sex. **C)** The correlation heatmap constructed by WGCNA showing relationship between identified modules and sample categories. Modules must contain >= 50 genes, otherwise are merged. **D)** Scatterplots of top highly correlated modules with age groups (3-19) in tumor or normal samples. R represents regression value of the corresponding line. The highlights show confidence of intervals. Colors of the dots, lines and highlights refers to the type of gene (Oncogene, TF, other). Abbreviations: Onco: oncogene, TF: transcription factor, TF/Onco: transcription factor/ oncogene, MM: module membership, GS: gene significance,

To analyze DEGs, samples from infants (0-2) and childhood (3-19) were compared separately for normal and tumor tissues. The differential expression status of each gene was then compared between the DEG lists of normal and tumor tissues to identify genes that exhibit significant changes in one of the tissues. Genes were considered age-specific if they met the following criteria: FDR ≤ 0.05, an average logCPM expression ≥ 0.5, and a log2 fold change ≥ 1.5 or ≤ −1.5 in one comparison (e.g., 0-2 vs. 3-19 in normal tissues), while the log2 fold change was < 0 or > 0 in the other comparison, respectively. A total of 59 age- and tissue-specific DEGs were identified (Supp. file 1 S7).

The Weighted Gene Co-expression Network Analysis (WGCNA) method identifies co-expressed genes and associates these modules with specified traits within samples [30]. The results of the WGCNA analyses revealed three modules (14, 19, 21) associated with tumor tissues and one module (20) associated with normal tissues at the childhood stage (Fig. 3C). Scatterplots showing the correlation between module membership and gene significance values indicate a strong correlation between the identified modules and the state of the samples (Fig. 3D and Supp. Fig 4A). Specifically, modules 19 and 20 are positively correlated with childhood AML, while modules 14 and 21 are negatively correlated. Highly correlated genes within each module were selected as age-associated if they exhibited significance and module membership values greater than 0.5. Using these criteria, 30 genes were identified as age-associated in pediatric AML (Supp. file 1 S7). In total, 89 genes were recognized as age-associated in pediatric AML and used for further analysis.

Additionally, network analysis approaches were employed to visualize highly connected genes and study their expression across different age groups. The top highly connected genes displayed limited differences between age groups, except for module 20, which showed a clear distinction between normal tissues of different age groups that was absent in the corresponding tumors. The network and a heatmap of median gene expression across different age groups and tissues are presented in Supp. Fig 3.

### Age-associated genes in pediatric AML

I further examined the 89 age-associated genes through clustering and functional enrichment analysis to uncover potential underlying mechanisms. Unsupervised clustering of these genes identified five clusters associated with either normal or tumor tissues across different age groups (Fig 4 A, Supp. File 1 S7). The optimum number of clusters was identified using elbow plot technique (Supp. Fig 4 B). For instance, cluster 1 contains genes that exhibit higher expression in normal childhood AML samples but show low expression in childhood tumor samples, indicating an age-related association that is disrupted in tumors (Fig. 4 A and B). Cluster 2 includes genes with lower expression in childhood tumors, while expression remains high in the corresponding normal tissues. Clusters 3 and 4 encompass genes with subtle expression trends related to aging (Fig. 4A-B, and Supp. Fig 1).

**Figure 4.**
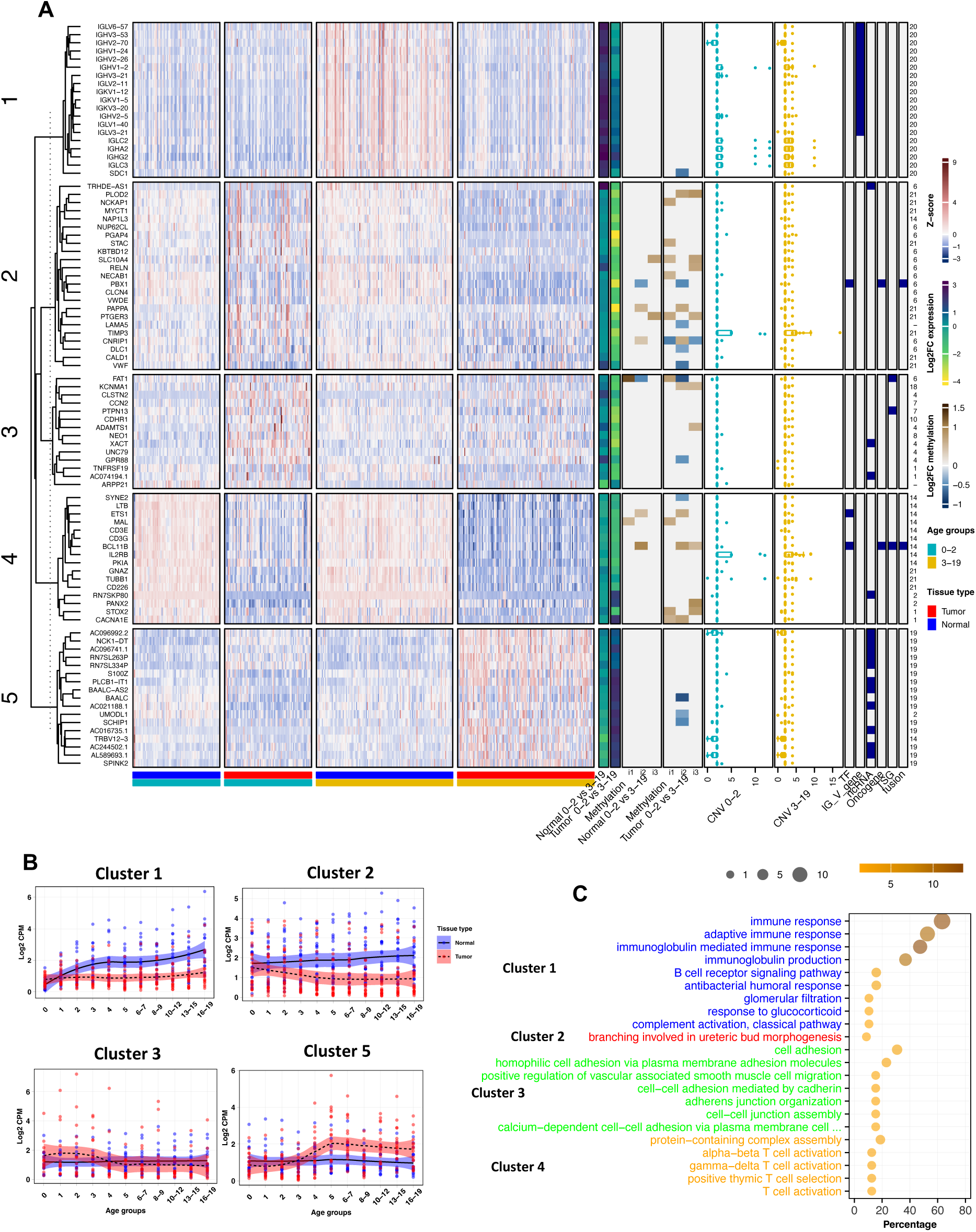
Integrated analysis of gene expression in infants and childhood. **A)** Heatmap showing expression of DEGs identified through comparisons of tumor and normal tissues in 0-2 and 3-19 groups. Each gene is associated with information: DEG status, differentially methylated regions (CpG islands, shores and shelves) between age groups, copy number variations, associated annotation, and number of modules identified by WGCNA. Genes (rows) are clustered according to Pearson correlation. i1) TSS, i2) gene body, i3) Other. **B)** The expression of genes within clusters undergoes significant changes over aging. In all clusters shifting in expression starts about year 2 and complete about year 4-5. Cluster 1, 2, and 5 show dramatic gene expression changes in tumors. In contrast, cluster 1 shows changes in normal tissues. Dots represent samples and the line show trend of changes. The highlighted sections represent confidence of intervals. **C)** Biological process associated to each module. Label colors represent different modules. Only up to 10 terms with a Pvalue <= 0.05 were included. Abbreviations: TF: transcription factor, TSG: tumor suppressor gene.

To investigate the epigenetic status of the age-associated genes, DMRs were analyzed in the following gene regions: Transcription Start Site (TSS) (i1), gene body (i2), and other regions (5’UTR, 1stExon, and 3’UTR) (i3) (Fig. 4A). The results indicated that DMRs are more frequently associated with tumors and remain unchanged in normal tissues. Notably, DMRs were more common in genes of Cluster 2, correlating with gene expression changes observed in this cluster (Fig. 4A). To assess the relationship between gene expression and genetic variation, CNV was analyzed in both age groups. The results showed no significant differences in CNV between the age groups (Fig. 4A). Interestingly, genes in Clusters 1 and 5 were identified as belonging to similar groups: immunoglobulin genes and long noncoding RNAs (lncRNAs), respectively. This suggests potential functional disruption by cancer in these gene groups (Fig. 4A).

To understand how age-associated gene expression changes over time, expression trends were analyzed annually up to 5 years of age and in 2-4 year bins up to 19 years. The trend lines for each cluster indicated that expression changes typically begin around age 1 and stabilize by ages 4-5 (Fig. 4B). Four distinct patterns of expression changes with aging were observed: (i) expression increases in normal tissues but remains unchanged in tumor tissues (Cluster 1), (ii) expression increases in tumors but remains stable in normal tissues (Cluster 5), (iii) expression slightly increases in normal tissues but decreases in tumor tissues (Cluster 2), and (iv) expression changes are minimal with aging (Clusters 3 and 4) (Fig. 4B, Supp. Fig 4C). Functional annotation revealed that Clusters 1, 3, and 4 are primarily associated with immune response and cell adhesion, while Cluster 5, which comprises mainly lncRNA-encoding genes, did not show specific functional annotations (Fig. 4C). Cluster 1 is related to adaptive and immunoglobulin-mediated immune responses, with numerous immunoglobulin genes involved (Fig. 4A and C). Cluster 2 is associated with ureteric bud morphogenesis, linked to LAMA5 and PBX1 genes (Fig. 4C). Cluster 3 is related to cell adhesion and cell-cell junctions, while Cluster 4 is associated with T cell activation, though genes in this cluster show only subtle changes with aging (Fig. 4C). Cluster 5, consisting of 11 lncRNA-encoding genes, lacks functional associations but demonstrates distinct expression patterns across different age groups, suggesting potentially important functions.

Altogether, 89 genes were introduced and analysed that show age associated expression pattern that change occur during the transition from infancy to childhood, specifically between ages 1 to 4. These genes are predominantly associated with immune response and cell adhesion functions.

### Transcriptional regulatory network conferring the infant and childhood AML

The distinctions between infant and childhood AML are potentially associated with a regulatory circuitry that underlies these differences. To identify age-associated TFs in pediatric AML, I employed the same approach used for identifying age-associated genes, but with less restrictive cut-offs. For differential expression analysis, a log2 fold change ≥ 1 and ≤ −1 was applied. For WGCNA-identified genes, significance and module membership values greater than 0.2 were used. Consequently, the number of identified TFs was larger than the previous comparisons (Fig 4), with a trade-off in specificity of expression (Fig. 5A). A total of 18 TFs associated with age groups of AML patients were identified that by overlaying these identified TFs on the TF-gene interaction regulatory map (hTFtarget) [34], a regulatory network of TFs was constructed (Fig. 5A, Suppl. File 1 S11-12). This network includes only TFs that function in the bone marrow, with non-TF genes excluded to focus on TF interactions and identify the most connected TFs within the regulatory network. Sorting TFs based on centrality factors, including degree and betweenness, revealed the central TFs of the network. The top five important TFs, which are also differentially expressed between age groups, are ZBTB46, PBX1, ETS1, TSC22D1, and ZNF888.

**Figure 5.**
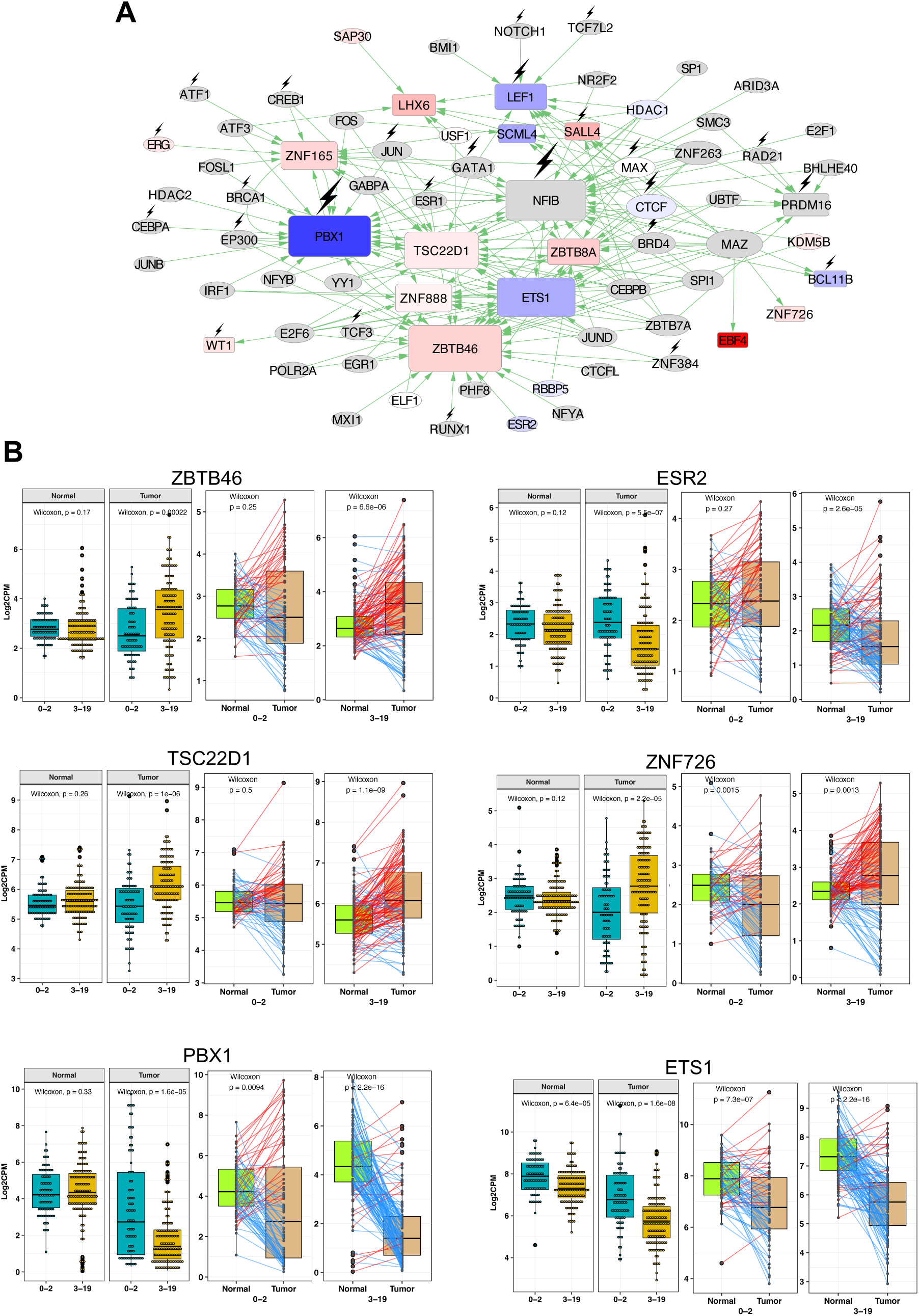
TF regulatory network of infant and childhood AML. **A)** Transcriptional regulatory network of infant and childhood AML. The regulatory network is constructed by mapping candidate TFs on hTFtarget network (Zhang,Q. et al. 2020). TFs were selected based on differentially expression and detection by WGCNA. The network was filtered for only interactions identified in bone, and bone marrow. Gray nodes are not differentially expressed. Square nodes are detected in this study and circular nodes are other associated TFs by hTFtarget. Larger nodes represent higher connectivity. Blue and red colors indicate the gradient of down and up regulated TFs, respectively, with darker color shows higher change in expression. Arrows show the direction of connections. Lightening sign indicates oncogene activity. **B)** Box plots showing the expression level of candidate TFs in infant and childhood samples. Comparisons are performed between different age groups of the same tissue type (left boxes) and different tissues in each age group (right boxes). The comparisons of different tissues were performed by Wilcoxon test between paired samples (connected by lines). Red and blue lines show increased and decreased expression, respectively, in same patients.

Visualization of differentially expressed (DE)-TFs showed age-specific expression patterns in tumor or normal tissues. For instance, ZBTB46, TSC22D1, and ZNF726 exhibit higher expression in tumors at the childhood stage compared to the infant stage, while ESR2 shows lower expression in childhood tumors (Fig. 5B). PBX1, ETS1, and LEF1 display reduced expression in tumors of both age groups, with a more pronounced reduction in childhood patients (Fig. 5B and Supp. Fig 5). Expression comparisons of other identified age-associated TFs are detailed in Suppl. File 1 S12. Many of these TFs show significant expression variation in tumor tissues, likely due to tumor diversity and mutations.

In summary, multiple age-associated TFs were identified using differential expression analysis and WGCNA. A transcriptional regulatory network was constructed based on these TFs, and the central TFs within the network were identified. Expression analysis of these TFs across different tissues and age groups revealed age-specific expression patterns.

### WGCNA identifies multiple gene groups associated to pediatric AML

The WGCNA identified several modules associated with pediatric AML (Fig. 3C, Supp. File 2). Eight modules were identified, with four (Modules 3, 4, 5, and 16) positively correlated with AML tumors and four (Modules 1, 2, 6, and 14) negatively correlated. Most analyses focused on the top six modules. Modules 3, 4, and 16, which are highly correlated with AML, encompass multiple TFs and oncogenes. In contrast, Modules 1, 2, and 6 are negatively correlated with the tumor and contain relatively fewer TFs and oncogenes (Fig. 6A-C, Supp. Fig 4A). Only highly correlated TFs and oncogenes with a gene significance value of ≥0.5 are shown for each module. The identified TFs in these modules exhibit high correlation in expression and may play significant roles in pediatric AML.

**Figure 6.**
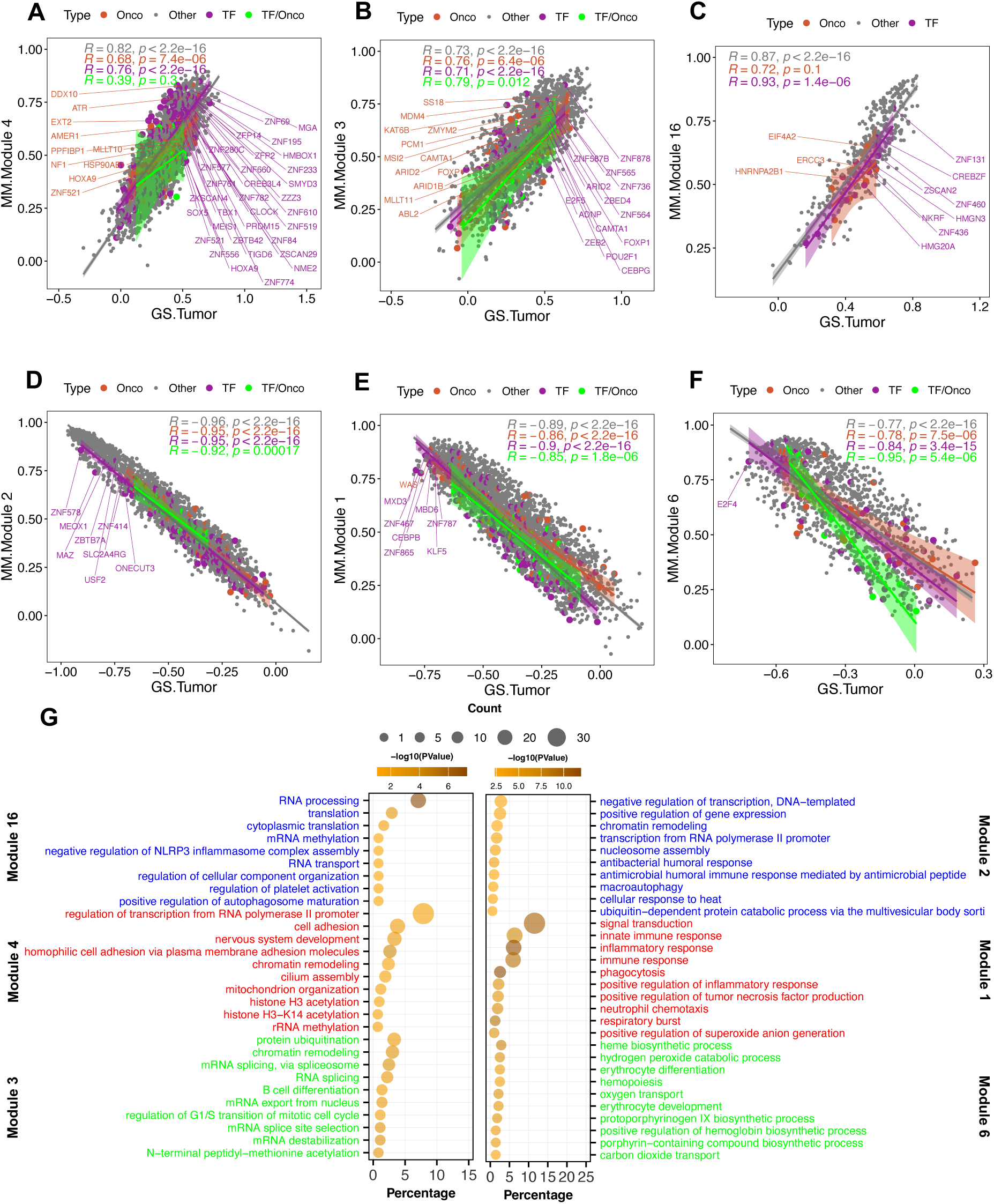
Scatter plot and gene ontology analysis of highly associated modules with AML. **A-C)** Correlated modules with tumor samples. **D-F)** Anti-correlated modules with tumor samples. In A-F, R represents regression value of the lines. The names of genes of interest with a gene significance value >= 0.5 are shown. Highlights show confidence of intervals. Colors of the dots, lines and highlights refers to the type of gene. **G)** Biological processes associated to modules. The label color represents different modules. Abbreviations: Onco: oncogene, TF: transcription factor, TF/Onco: transcription factor/ oncogene, MM: module membership, GS: gene significance. Count: number of genes out of the total submitted with a hit for annotation. Percentage: percentage of genes out of the total submitted with a hit for annotation.

Functional annotations of the modules reveal that positively correlated genes are generally associated with transcription, RNA processing, and development, while negatively correlated modules are primarily related to transcription, immune response, and hematopoiesis (Fig. 6G). Modules 16 and 3 are mainly associated with RNA processing, translation, and RNA splicing. Members of Module 3, including SMARCE1, CDKN2B, SMARCC1, PHF10, and CDKN2A, are also linked to cell cycle development. Module 4 is associated with transcription and chromatin remodeling. Module 2 is related to transcription, chromatin remodeling, and immune responses. Module 1 is involved in various immune response processes, including inflammatory response, phagocytosis, and neutrophil chemotaxis. Module 6 is associated with hematopoiesis, erythrocyte differentiation, and development (Fig. 6G). These findings suggest that the modules identified by WGCNA are linked to processes involved in gene regulation, immune response, and blood cell development, all of which are pertinent to pediatric AML.

Network based analysis of the modules identified most central genes in the network based on the weighted degree analysis (Supp. Fig 6). For each module, top 10 most connected genes, the centrality degree and the differentially expressed gene status is shown as bar plots (Fig 7 A and B). Some of most central genes in the networks, for example AC023818.1 from module 3, are not DEGs, indicating that the gene is selected based on connection with many numbers of genes within the module. In addition, many of the identified genes, particularly within module 1, 2, 3, and 16, encode non-coding RNAs or pseudogenes that yet might be important for the AML cancer diagnostics or developing new therapeutic strategies.

**Figure 7.**
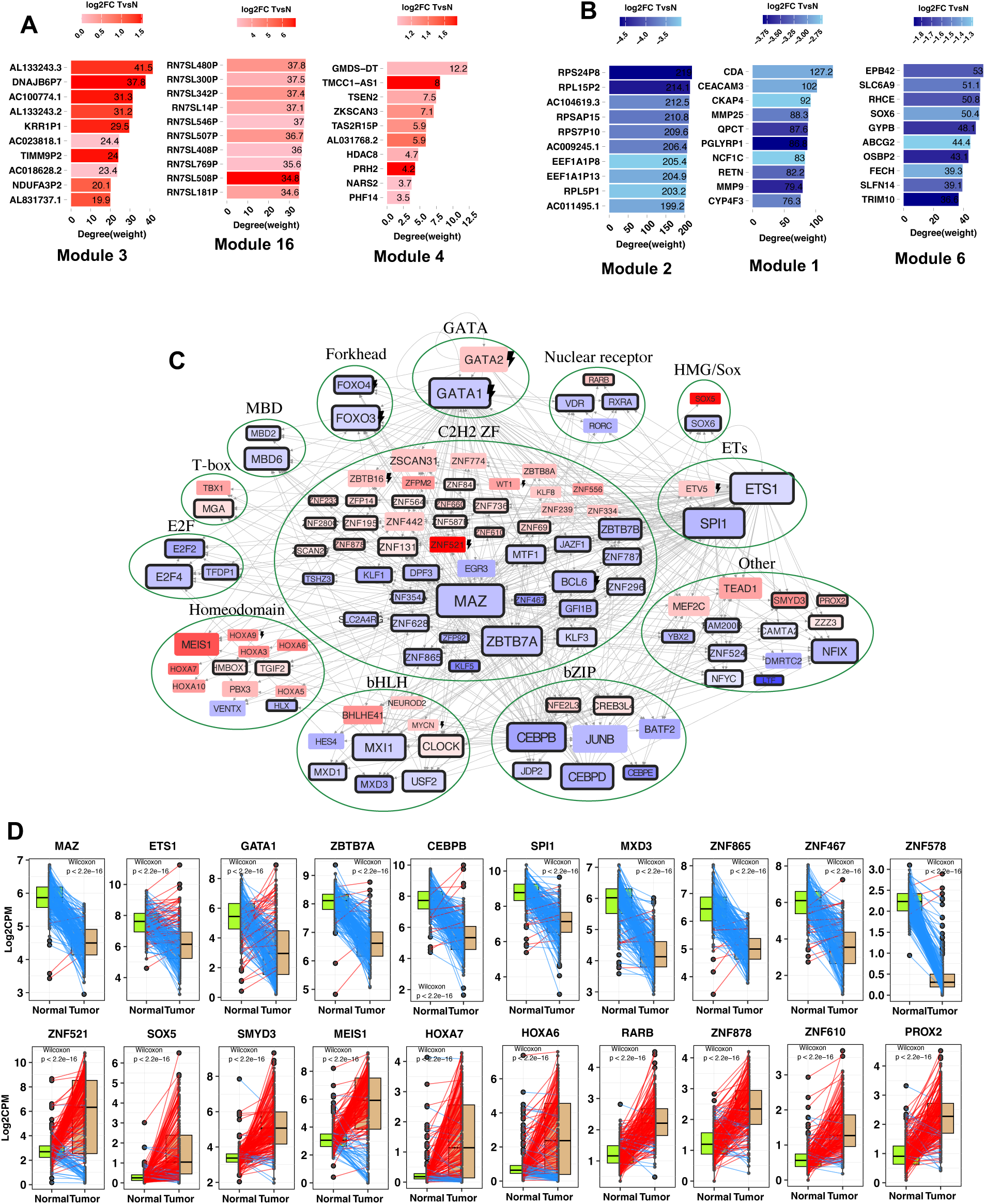
Candidate genes and TF regulatory of pediatric AML. **A and B)** Bar plots show top 10 genes and log2fold change of tumor vs normal tissues. **C)** Transcriptional regulatory network controlling pediatric AML. The network is constructed by mapping highly correlated TFs identified by WGCNA and differentially expressed TFs on hTFtarget network (Zhang,Q. et al. 2020). Star sign shows significantly differentially expressed TFs with FDR <= 0.05 and Log2FC ± 0.585. Lightening sign indicates oncogene activity. The purple border of nodes indicates detection by WGCNA. Arrows head shows the regulatory direction between nodes. Larger nodes represent higher connectivity. Blue and red colors indicate the gradient of down and up regulated TFs, respectively, with darker color shows higher change in expression. Nodes are grouped based on the family of binding domain. **D)** Bar charts show gene expression level of top 10 highly connected TFs and highly differentially up- or down-regulated genes. The comparisons were performed by Wilcoxon test between paired samples (connected by lines). Red and blue lines show increased and decreased expression, respectively, in same patients.

In summary, WGCNA identified several modules highly correlated with pediatric AML tumors. These modules include multiple transcription factors (TFs) and oncogenes, which may play significant roles in cancer regulation. The modules are associated with biological processes such as transcription regulation, immune response, and blood cell development, all of which are relevant to AML. Additionally, we have identified several novel markers that could potentially be used for personalized medicine and therapeutic purposes.

### A potential regulatory network for regulating pediatric AML

The extensive changes in gene expression observed in pediatric AML suggest alterations in gene regulatory mechanisms. To explore these mechanisms, I constructed a regulatory network using TFs identified through differential expression analysis and WGCNA (Fig. 7B). TFs were considered differentially expressed if they met the following criteria: FDR ≤ 0.05, log2 average CPM ≥ 0.5, and log2 fold change ≥ 1.5 or ≤ −1.5. These cut-offs ensure the selection of rigorously differentially expressed TFs. For TFs identified by WGCNA, I selected those from eight highly associated modules (1, 2, 3, 4, 5, 6, 14, and 16) with module membership values ≥ 0.7 and gene significance values ≥ 0.5. The resulting list of TFs was then mapped onto the hTFtarget [34] database, filtered for interactions within the bone marrow. The final network comprised 463 edges and 109 nodes (Suppl. File 1 S9-10).

Network analysis revealed the most connected TFs within the network. The top 10 most connected TFs were MAZ, ETS1, GATA1, SPI1, ZBTB7A, CEBPB, MXI1, JUNB, CEBPD, and FOXO3. All these TFs, except JUNB, were identified through WGCNA analysis and are differentially expressed. Notably, SPI1, CEBPB, JUNB, and CEBPD exhibited a log2 fold change < −1.5. Additionally, TFs were clustered based on their DNA binding domains [35] to illustrate their involvement in pediatric AML. The analysis showed that members of the C2H2 zinc finger family, followed by homeodomain, bHLH (basic helix-loop-helix), and bZIP (basic leucine zipper) families, are more frequently involved in pediatric AML. The expression patterns reveal that C2H2 zinc finger family members are almost equally up- and down-regulated in tumor samples, whereas homeodomain family members are predominantly up-regulated and bZIP members are mostly down-regulated (Fig. 7B). In smaller families within the network, down-regulation is more frequent.

The connectivity within the network is influenced by the availability of ChIP-seq information. To further explore gene expression patterns, I visualized the expression of the top 5 highly differentially expressed genes within the network, irrespective of their connectivity status (Fig. 7C). Box plots of paired samples (from the same patients) reveal dramatic and significant changes in the expression of these TFs in pediatric AML tumors (Fig. 7C).

Altogether, I have developed a regulatory network for pediatric AML based on differential expression analysis, WGCNA outputs, and available protein-gene interaction data. This network highlights multiple highly connected and variable transcription factors (TFs) that could be valuable for further cancer research.

### Novel markers for pediatric AML

To summarize the novel findings of this study, Table 1 presents the genes identified from various analyses. This table categorizes the genes based on their method of identification and current knowledge. These genes have potential applications in developing new markers for infant, childhood, and generally pediatric AML. Additionally, the identified genes and pathways may provide new insights for developing innovative therapeutic approaches for pediatric AML.

**Table 1.**
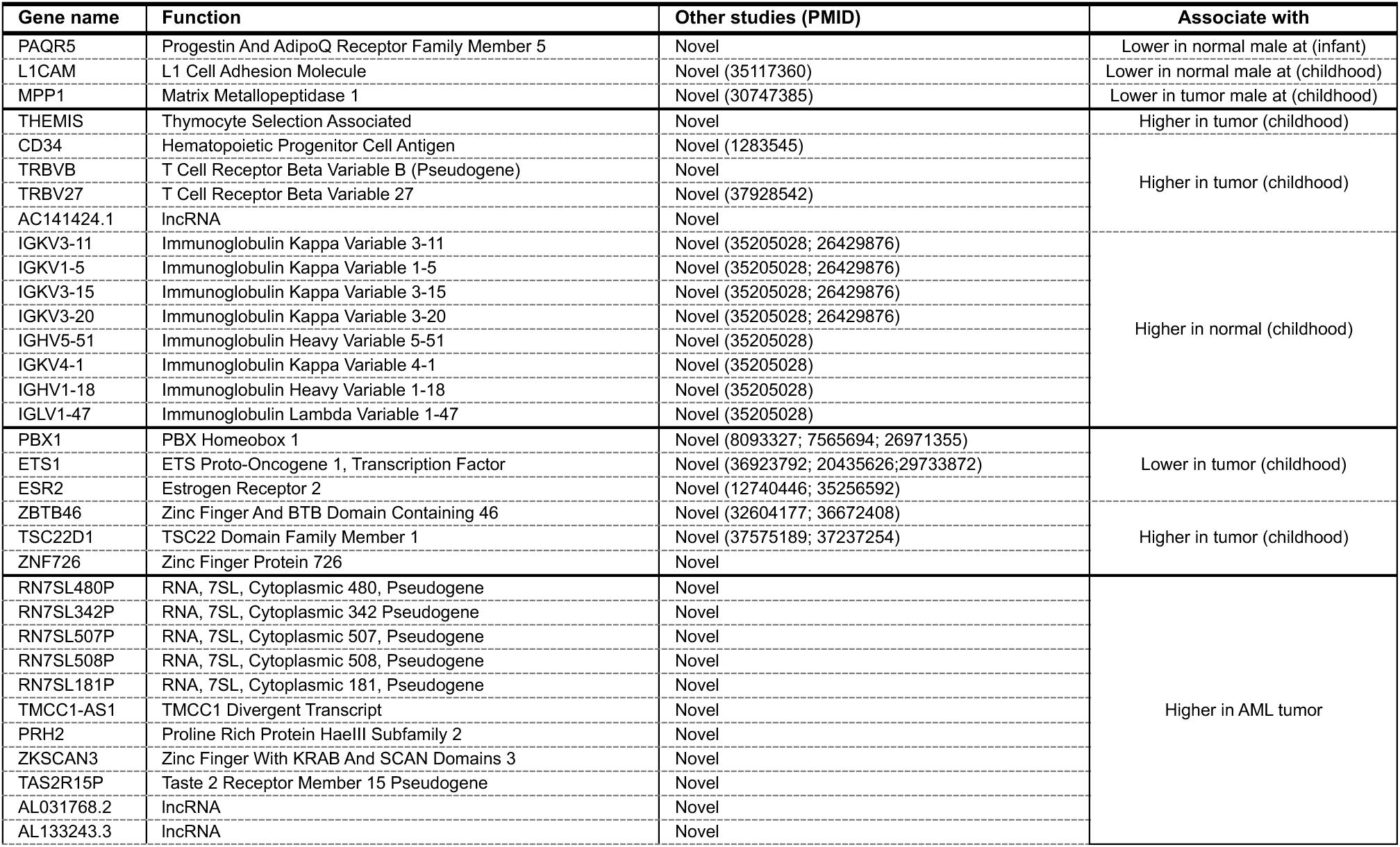
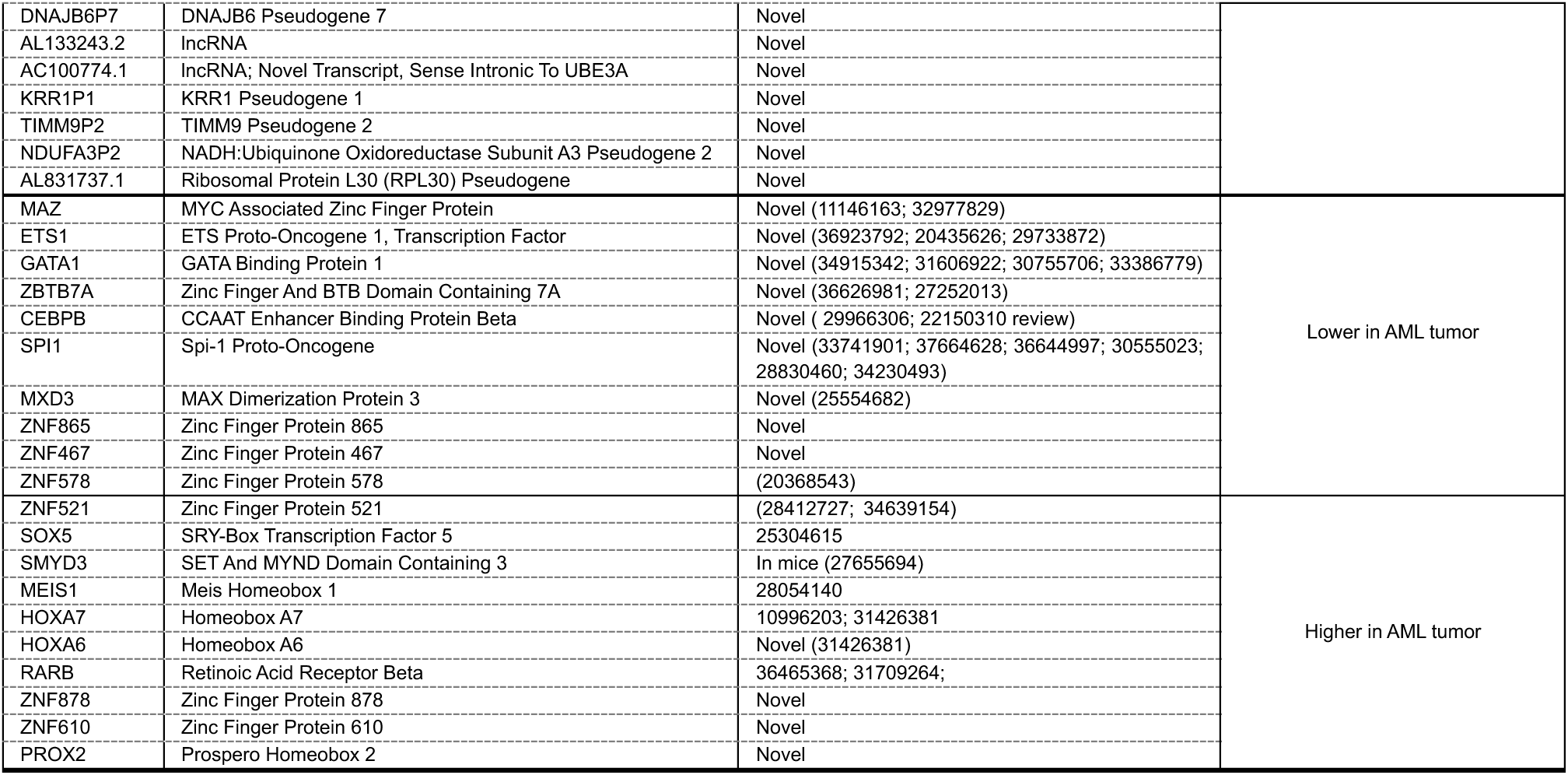
Remarkable identified genes through network and differential expression analysis. If the identified gene is associated to AML in a publication, the corresponding PMID is shown.

## Discussion

Pediatric AML is a heterogeneous disease, categorized based on clinical features, cytogenetic, and molecular aberrations. Significant heterogeneity is observed between infant and childhood AML patients, with distinct cytogenetic abnormalities [11, 14], mutations [14–19], and molecular pathways aberrations [14, 20]. While some high-throughput studies have revealed transcriptional differences between pediatric and adult AML [21, 22], and network-based analyses have identified subtypes, regulatory networks, hub genes, and pathways in AML [36–39], no studies have specifically compared the transcriptomes of infant and childhood AML patients. In this study, I utilized transcriptome data from 406 paired normal and tumor tissues, along with graph-based analysis, to investigate the transcriptional distinctions and gene regulatory mechanisms between infant and childhood AML. Additionally, I present a comprehensive regulatory network for pediatric AML.

This study utilized paired primary AML tumor samples collected from bone marrow tissues of patients under 20 years of age, resulting in a closely matched cohort. To maximize insights from the data, all types of genes—protein-coding and non-coding, including lncRNAs and pseudogenes—were included in the analysis. The role of lncRNAs [40, 41], and pseudogenes in cancer is well-documented [42, 43]. For differential expression analysis, the Wilcoxon rank-sum test was employed, as conventional methods such as edgeR and DESeq2 tend to produce an exaggerated number of DEGs due to numerous false positives [26]. This method has been successfully used in other studies and single-cell RNA sequencing analyses involving large sample sizes [44–48]. The use of the Wilcoxon rank-sum test resulted in a more accurate number of DEGs, which were subsequently validated through heatmaps and boxplots. Additionally, applying rigorous statistical thresholds (FDR, log2FC, and expression levels) further refined DEG identification, ensuring that the selected genes exhibited robust expression alterations.

Correlations between genes and their associations with sample characteristics can reveal underlying biological mechanisms. In this study, WGCNA was utilized to exploit the intrinsic characteristics of the data to identify gene associations with infant and childhood age groups in AML [30]. WGCNA successfully identified gene modules associated with these age groups and specific tissues. The analysis revealed multiple gene groups positively or negatively correlated with pediatric AML. Modules, as subnetworks, were analyzed to detect central genes that might play critical roles due to their extensive connections within the network [32, 49]. The expression of highly connected genes within these modules suggests their involvement in distinct biological pathways [50]. The identified modules related to AML are primarily associated with immune response, erythrocyte development, and gene regulation. Notably, the results show a down-regulation of genes linked to immune response, a finding consistent with observations in both pediatric and adult AML [51]. Furthermore, the analysis highlights the importance of transcriptional regulation in cancer development, emphasizing the critical role of transcription factors (TFs) in pediatric AML [52, 53].

Despite the phenotypical distinctions between infant and childhood AML, there is limited knowledge about their transcriptional differences. By combining differential expression analysis and WGCNA, I identified genes associated with age groups that exhibit varying expression levels between normal and tumor tissues. Generally, the differences in expression patterns emerge around age 1 and are well-established by ages 4-5. This temporal pattern aligns with the phenotypical differences observed between infant and childhood AML [11, 14–20]. The results indicate that genes related to immune response, particularly those encoding immunoglobulins, increase in normal tissues but remain unchanged in tumors. Although immunoglobulin levels rise with age in infants [54, 55], pediatric AML tumors, whether from infants or childhood patients, show similar levels of immunoglobulins. It has been shown that dysfunctional antibodies promote tumor progress [56, 57]. This observation suggests that low levels of immunoglobulins may be necessary for AML progression, which could be exacerbated in infants due to their inherently immature immune systems. Cell adhesion is only subtly affected across different age groups. Dysregulation of cell adhesion in AML is associated with several processes, including migration and quiescence [58]. Additionally, multiple age-associated lncRNAs were identified, showing dramatic changes in tumors with aging. The role of lncRNAs in AML has been well-documented [59, 60], suggesting they may play significant roles in distinguishing between infant and childhood AML.

I constructed a potential regulatory network to explore the distinctions between infant and childhood AML. As anticipated, only a few differentially expressed transcription factors (DE-TFs) were identified, consistent with the relatively small number of age-associated differentially expressed genes (DEGs). The changes in TF expression levels are subtle. For instance, genes like ETS1 and PBX1 exhibit increased expression in one age group, suggesting that minor variations in TF expression could contribute to the differences observed between infant and childhood AML. Network analysis highlighted ZBTB46 as a crucial TF, showing higher expression levels in childhood AML patients. ZBTB46 has been previously reported to be upregulated in AML tumors [61]. PBX1, another significant TF, is known for its role in pediatric acute lymphoid leukemia through its fusion with E2A [62]. In our study, PBX1 expression is notably low in infant tumors and decreases further in childhood AML, indicating its potential role in distinguishing between infant and childhood AML.

In this study, I analyzed pediatric AML samples to identify novel markers and therapeutic targets. Using WGCNA, several gene modules associated with pediatric AML were discovered. Generally, up-regulated modules are linked to gene regulation, RNA processing, cell adhesion, and development, while down-regulated modules are associated with immune response, transcription, and erythrocyte development. The significance of immune response in AML is well-documented [56, 57]. and erythrocyte development is relevant as AML impacts myeloid precursor cells that contribute to white blood cells, erythrocytes, and platelets [63]. Within each module, the most connected genes were highlighted as potential new markers or targets for AML diagnosis and treatment. These genes show high correlation with other genes and exhibit significant differential expression compared to normal tissues—key attributes for identifying cancer-associated genes [32, 49, 50]. Additionally, I constructed a regulatory network using differentially expressed TFs in pediatric AML. This network illustrates how TFs are interconnected based on protein-gene interaction data from numerous ChIP-seq analyses performed on bone marrow samples [34]. The network includes TFs from various families, reflecting extensive changes in gene expression in pediatric AML. Paired sample analysis revealed significant differences in TF expression across many patients, underscoring the potential importance of these TFs. While this network provides new insights into pediatric AML mechanisms, it also reaffirms the roles of previously identified TFs in this disease.

In conclusion, this study provides a comprehensive analysis of pediatric AML to reveal transcriptional distinctions between infants and childhood patients, regardless genomic abortions of patients. By identifying gene clusters associated with age and tissue type, and constructing a regulatory network, I have elucidated potential mechanisms underlying this heterogeneity. Additionally, a regulatory network of highly differentially expressed TFs specific to pediatric AML has been proposed. These findings highlight several new genes that could serve as a foundation for future research, including disease modeling, diagnostics, and therapeutic development.

## Acknowledgment

The results published here are in whole or part based upon data generated by the Therapeutically Applicable Research to Generate Effective Treatments (TARGET) initiative, phs000218, managed by the NCI. The data used for this analysis are available [fill in with NCBI dbGaP/SRA and/or NCI GDC/DCC url references]. Information about TARGET can be found at http://ocg.cancer.gov/programs/target. Toward a Shared Vision for Cancer Genomic Data [23]. R.G. did not receive any specific funding from any sources.

## Competing interest

The author declares no competing interests.

## Notes

### Competing Interest Statement

The authors have declared no competing interest.

