## Supplementary material for "Dissecting the transcriptional regulation of infant and childhood acute myeloid leukemia": Supplementary Figures.pdf

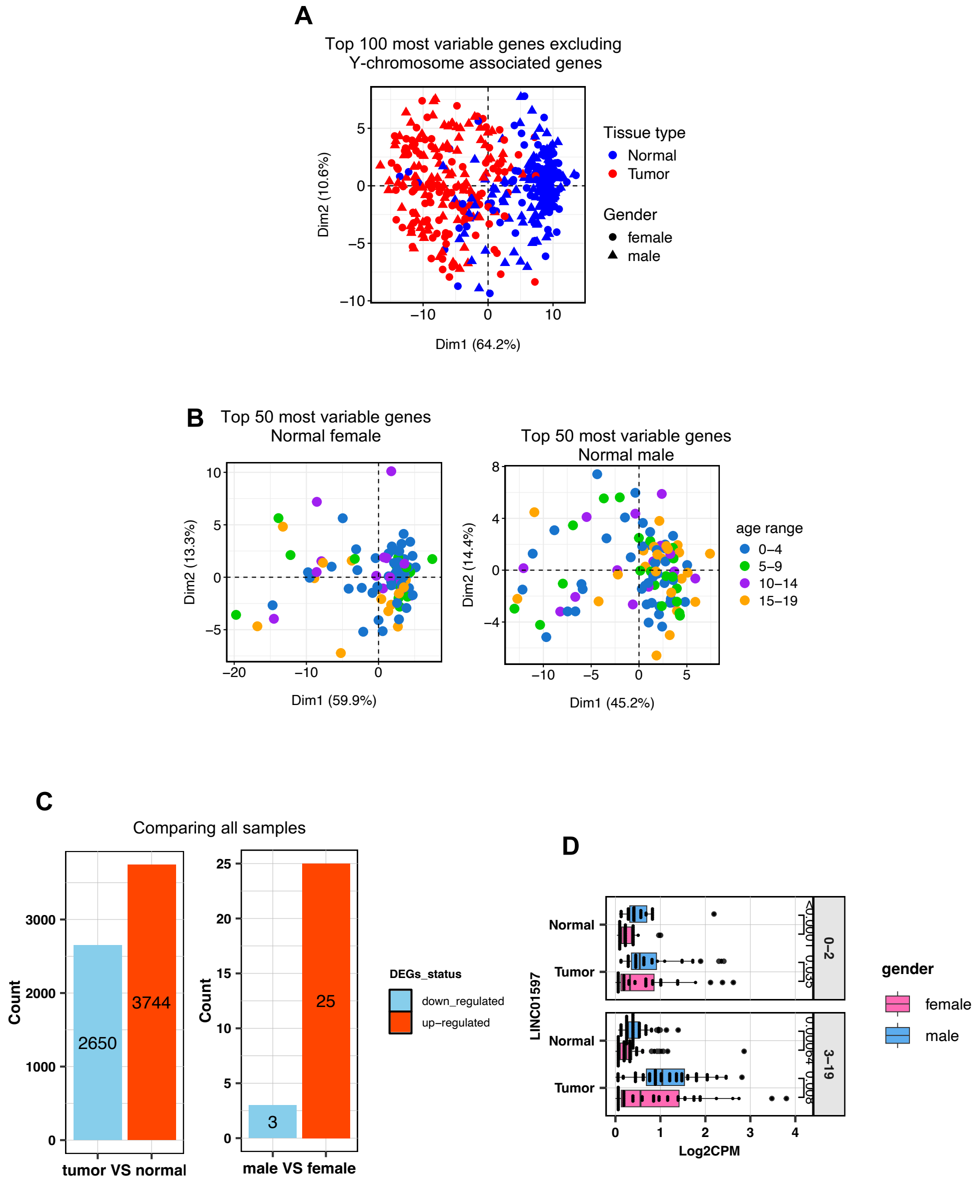


**Supplementary Figure 1**.

**A)** PCA of top 100 genes following removing Y-chromosome associated genes.

**B)** PCA plots of top 50 most variable genes in normal tissues of male and female patients does not show clustering based on age group.

**C)** Number of differentially expressed genes of comparison of all tumor vs all normal cells, and all male vs all female samples (regardless of tissue type).

**D)** Expression of LINC01597 in different tissues, sexes and age groups. LINC01597 show a relatively low expression. Wilcoxon rank-sum test was used to compare the means.

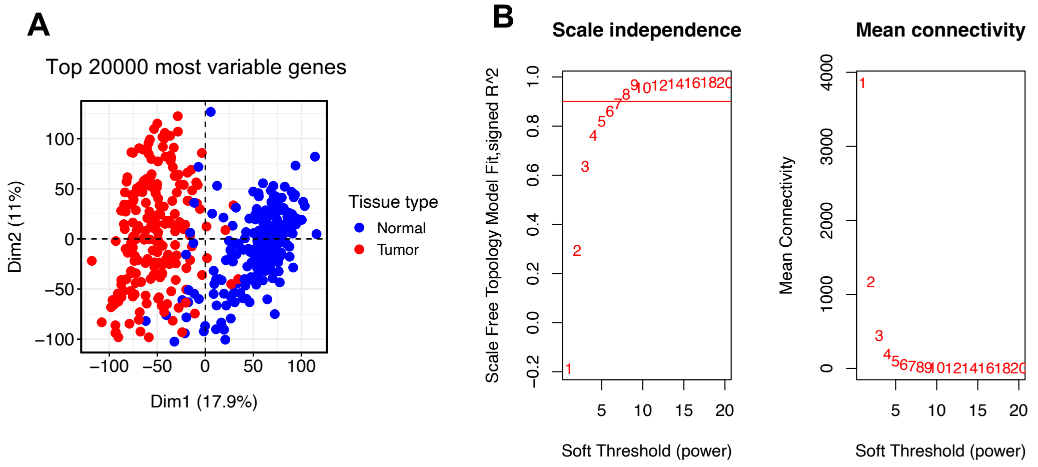


**Supplementary Figure 1**.

**A)** PCA plot showing cluster of samplings based on top 20000 most variable genes that were used for WGCNA.

**B)** Scale independence and mean connectivity plots showing soft power scores from 1 to 20.

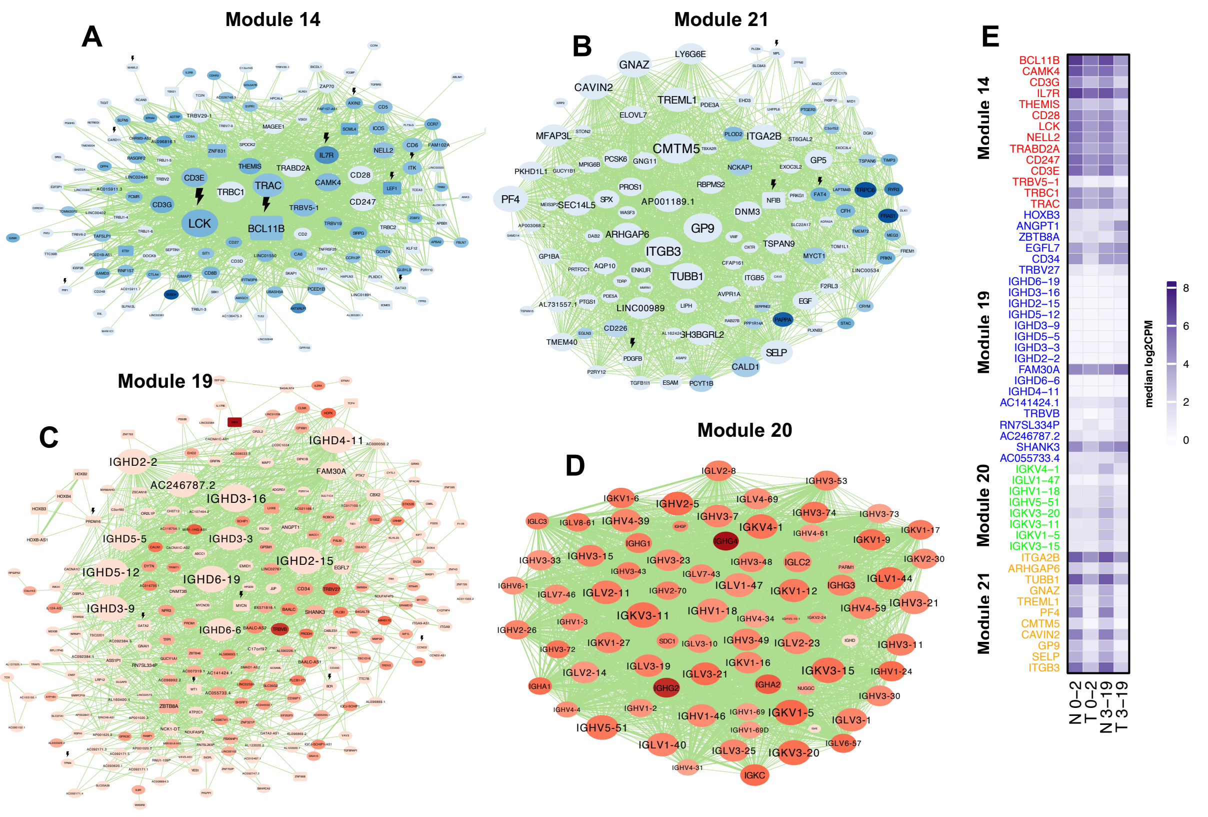


**Supplementary Figure 3**. Hub genes of the associated modules and the TF regulatory network of infant and childhood AML.

**A-D)** Networks showing modules highly associated with infant and childhood AML. Larger nodes represent hub genes. Darker blue and red colors represent down and up regulated genes, respectively, between age groups. Nodes with fade colors represent no significant differentially expression. Larger nodes represent higher connectivity. Blue and red colors indicate the gradient of down and up regulated TFs, respectively, with darker color shows higher change in expression. Thicker edges represented stronger connections. Lightening sign indicates oncogene activity.

**E)** Heatmap showing median gene expression of hub genes from networks in different normal and tumor tissues in infant (0-2) and childhood (3-19). Only top 10% highly connected genes (weighted Degree) are shown.

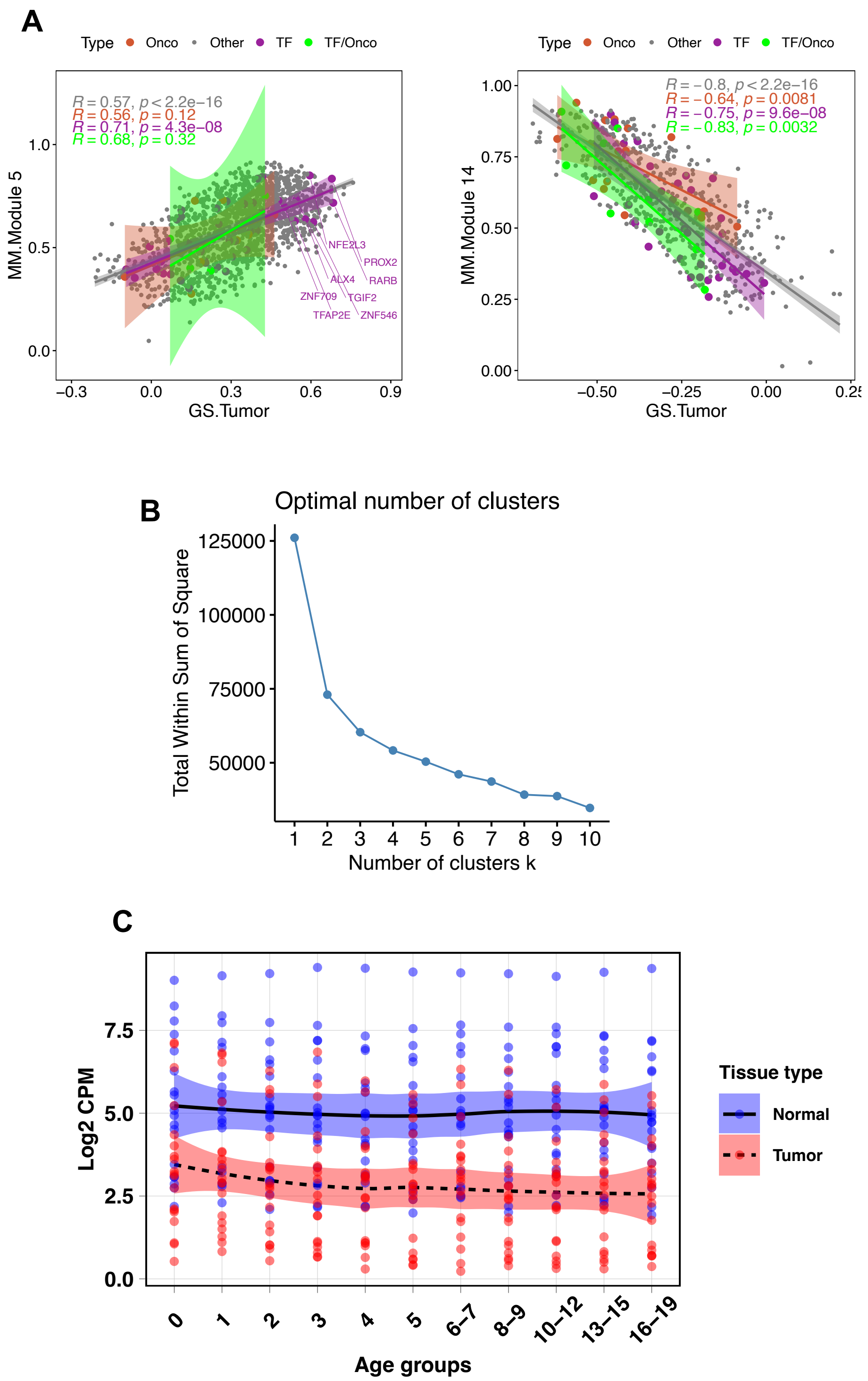


**Supplementary Figure 4.**

**A)** Scatterplots of module 5 and 14. associated to tumor in pediatric AML. R represents regression value of the corresponding line. The highlights show confidence of intervals. Colors of the dots, lines and highlights refers to the type of gene (Oncogene, TF, other).

**B)** Elbow plot showing the number of clusters for 89 age associated genes.

**C)** Cluster 4 of age group associated genes in pediatric AML.

Abbreviations: Onco: oncogene, TF: transcription factor, TF/Onco: transcription factor/ oncogene, MM: module membership, GS: gene significance.

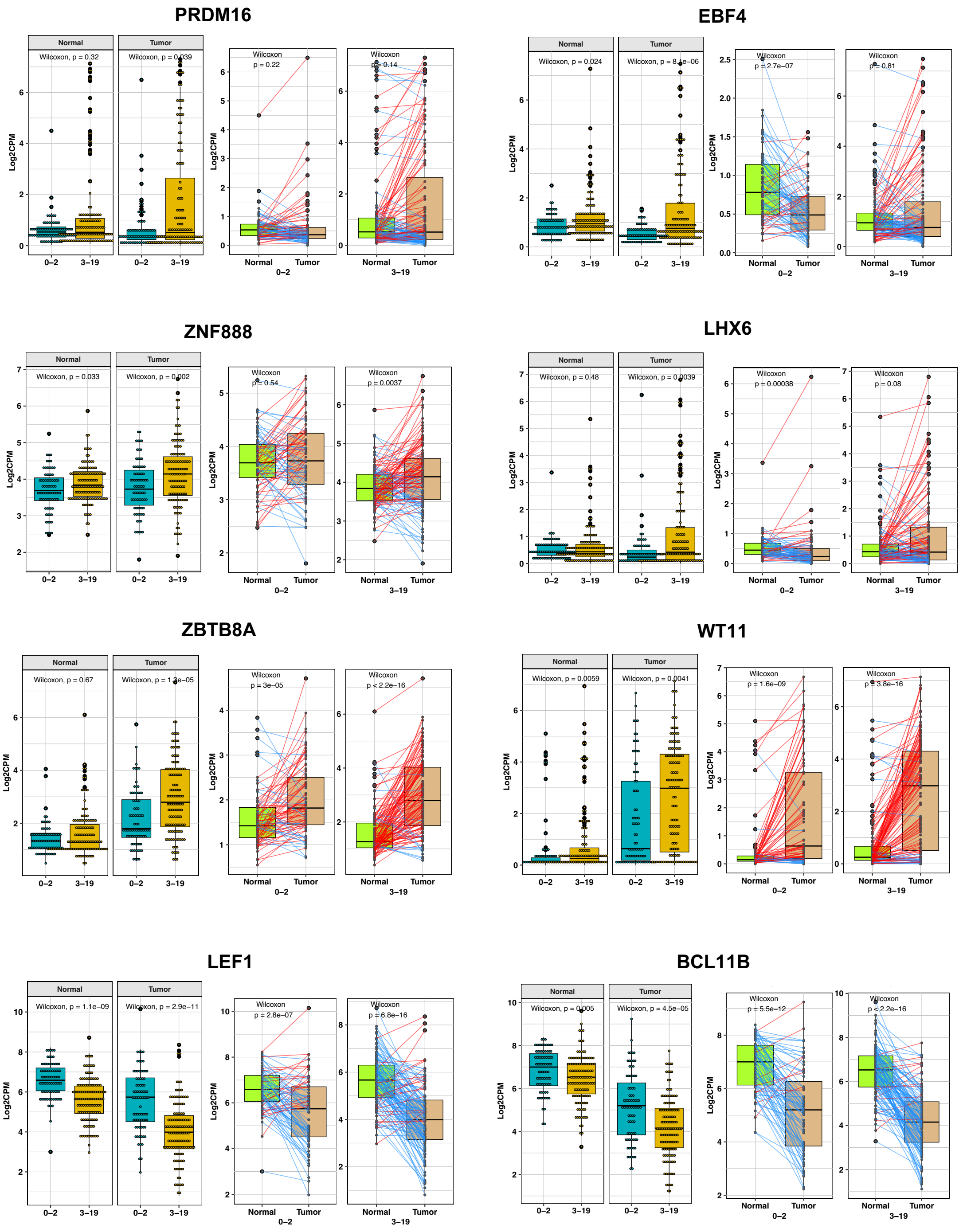


### Supplementary Figure 5.

Box plots showing expression level of transcription factors of infant regulatory network. For paired samples Wilcoxon signed rank test and unpaired samples Wilcoxon rank-sum test were used. Blue and red lines show lower and higher expression in the tumors, respectively.

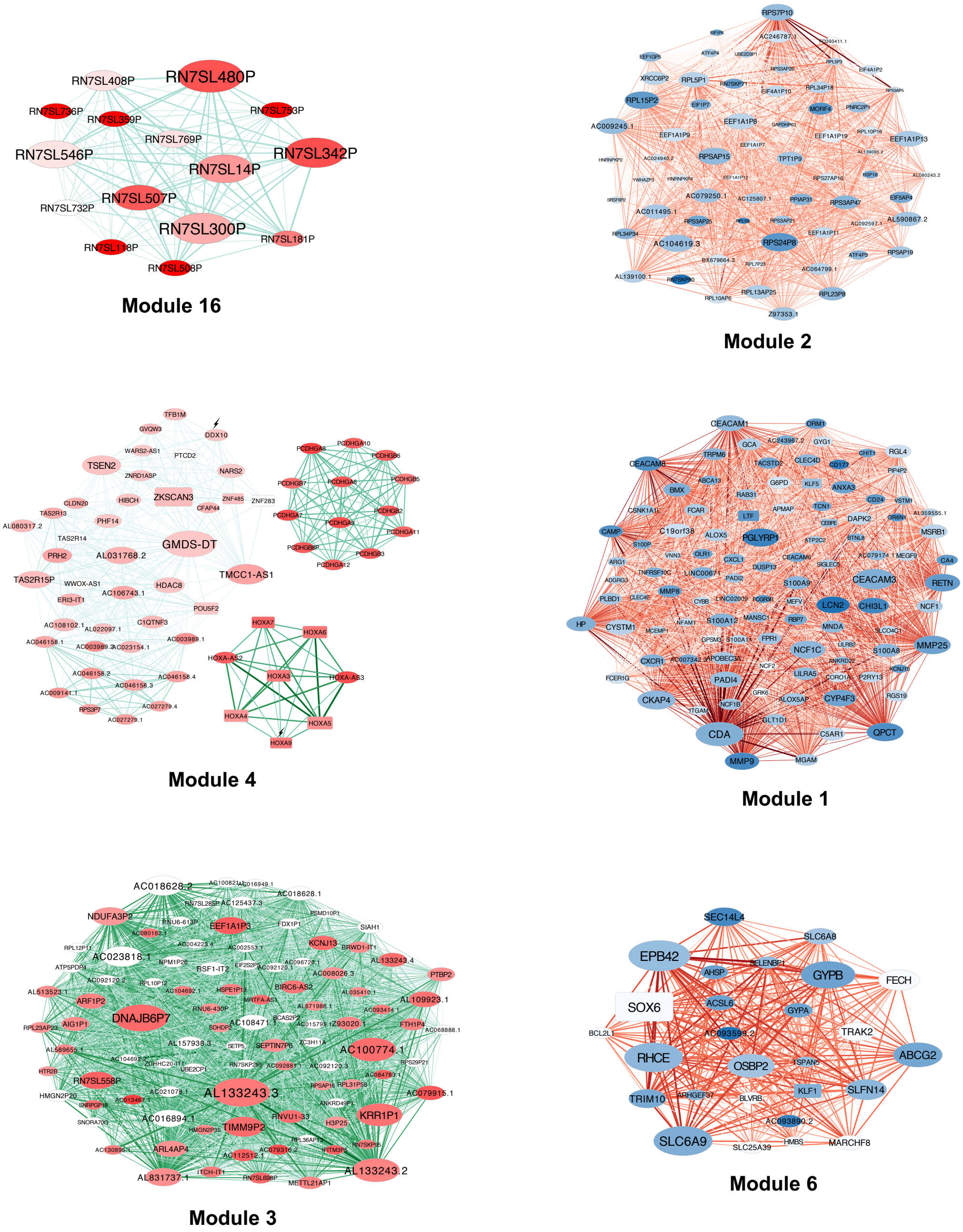


**Figure 6.** Gene networks of candidate modules associated to pediatric AML.

**A-F)** Networks of top 10% of highly connected nodes (based on weighted degree) in candidate modules. For each module top 10 genes and log2fold change of tumor vs normal tissues are shown in bar charts. Numbers on the bars show the weighted degree value. Larger nodes represent higher connectivity. Darker and wider edges indicate higher weights. Blue and red colors indicate the gradient of down and up regulated TFs, respectively, with darker color shows higher change in expression. White nodes represent not differentially expression.
